# The ectodermal loss of ARHGAP29 alters epithelial morphology and organization and disrupts murine palatal development

**DOI:** 10.1101/2025.03.11.642653

**Authors:** Emily Adelizzi, Lindsey Rhea, Campbell Mitvalsky, Samuel Pek, Bethany Doolittle, Martine Dunnwald

## Abstract

Orofacial clefts, including cleft palate (CP), are among the most common types of birth defects. CP specifically, results from a failure of palatal shelf fusion during development. Previous studies have shown that mutations in *RhoA GTPase Activating Protein 29* (*ARHGAP29)* are linked to CP, yet the role and tissue-specific requirements for ARHGAP29 during palatogenesis remain unknown. Here, we use tissue-specific deletion of *Arhgap29* in mice to provide the first direct evidence that ARHGAP29 is essential for proper palatal elevation and fusion. We demonstrate that ectodermal conditional loss of *Arhgap29* induces a significant delay in the fusion of palatal shelves at embryonic (E) day 14.5 and an incomplete yet significantly penetrant cleft palate at E18.5 – neither of which are observed when *Arhgap29* is lost later in development using K14-Cre. Phenotypic analyses of palatal shelves at E14.5 reveal a disorganized and thicker epithelium at the tip of the shelves. Loss of *Arhgap29* increases palate epithelial cell area and upregulates alpha-smooth muscle actin and phospho-myosin regulatory light chain implicating cell morphology and contractility as drivers of CP.

**Summary statement:** This study in mice is the first direct evidence that ARHGAP29 is essential for proper palatal elevation and fusion. Loss of *Arhgap29* alters oral epithelial morphology and upregulates contractility proteins.

## Introduction

Orofacial clefts (OFCs) are among the most common type of birth defects in humans, affecting roughly 1 in every 700 live births [1]. One type of OFC, non-syndromic (NS) cleft lip with or without cleft palate (CL/P), exhibits a complex etiology with both genetic and environmental underpinnings contributing to the defect [2]. One particularly well-defined gene regulatory network contributing to NSCL/P is the Interferon Regulatory Factor 6 (IRF6) gene regulatory network [3]. Mutations and single-nucleotide polymorphisms in *IRF6* cause and are associated with syndromic and NSCL/P, respectively [4, 5]. In addition to this direct role for *IRF6* in CL/P, genome-wide association and exome sequencing studies have revealed that mutations in downstream effector genes, such as *RhoA GTPase Activating Protein 29* (*ARHGAP29)*, are associated with and causative of CL/P, yet these studies are lacking functional evidence for a role of ARHGAP29 in CL/P [3, 6–8].

Palatogenesis is the embryonic process by which the palate forms. Starting at E12.5 in the mouse, the walls of the maxillary processes undergo bilateral growth, giving rise to two palatal shelves. Palatal shelves are comprised of a mesenchymal core containing neural-crest-derived mesenchymal cells underlying a layer of ectoderm-derived oral epithelial cells, themselves covered by a single layer of specialized squamous epithelial cells termed periderm [9, 10]. These palatal shelves grow vertically alongside the tongue (E13.5) and remodel to a horizontal position above the tongue by E14.0. They subsequently grow towards each other and the ectoderm-derived cells (forming the so-called medial edge epithelium, MEE) covering the mesenchyme begin undergoing major morphogenic events; first they make contact, then the superficial periderm is removed, followed by the intercalation of epithelial cells into a single cell layer (called the medial epithelial seam, E14.5), and finally they are removed as the palate forms (E15.0). This removal leads to the formation of a confluent mesenchymal bridge on the roof of the mouth formally giving rise to the separate oral and nasal cavities [11–14]. When the palatal shelves fail to fuse, this results in a cleft palate, which is lethal in the mouse as the animals cannot suckle and feed at birth. Loss of many genes causes cleft palate in the mouse, including those coding for transcription factors (i.e. IRF6, P63) [15, 16], growth factors (i.e. TGFb3, FGF10)[17–19], cell adhesion proteins (i.e. E-cadherin, Nectin-4) [20–22], and cytoskeleton (i.e. SPECC1L, β-catenin) [23–25], many of which belong to or are influenced by the RhoA pathway, suggesting that cytoskeletal remodeling may be driving palatogenesis.

RhoA is a small GTPase that shuttles between an active and inactive form based on its binding to GTP or GDP, respectively. Guanine nucleotide exchange factors (GEFs), Guanine nucleotide dissociation factors (GDIs) and GTPase activating proteins (GAPs) regulate this activity [26, 27]. Over 60 GAPs have been identified, and among them, the PTPL1-associated RhoGAP1 (PARG1, reclassified as ARHGAP29) was described as a specific inhibitor of RhoA signaling with a strong expression in human heart and skeletal muscle tissues [28]. Subsequent studies found ARHGAP29 in many other human and murine tissues and cell types, including endothelial cells, keratinocytes, and to a lesser degree fibroblasts [3]. Most of the initial work on ARHGAP29 was done in endothelial cells, where it reduces RhoA activity at cell-cell adhesion sites to promote their remodeling during tubulogenesis [29–31]. Subsequent studies have found that ARHGAP29 contributes to the definition of apical-lateral border polarity in epithelial tight junctions [32] and participate in the specification of the non-neuronal ectoderm fate [33]. In cutaneous keratinocytes, it is required for cell shape, proliferation, and migration [34] while during cancer metastasis, ARHGAP29 is upregulated and mediates Yes-Associated Protein 1 (Yap)-dependent actin remodeling in response to tissue stiffness [35–37]. Collectively, these studies demonstrate the importance of ARHGAP29 in various developmental events. This is further supported by the findings that *Arhgap29^fl^*;*Sox2-Cre* and *Arhgap29*-deficient murine embryos die prior to chorioallantoic fusion (before E8.5) [29, 38, 39]. We previously identified several *ARHGAP29* genetic variants in patients with CL/P [3, 40] and subsequently developed a murine model with a point mutation that mimics an *ARHGAP29* variant (K326X) found in a patient with CL/P [38]. This single-nucleotide, nonsense mutation produces a truncated protein that is presumed to lack function because no homozygous *Arhgap29^K326X^* offspring were observed (as early as E8.5). Due to this early lethality of the global *Arhgap29* knockout, this experiment did not shed light on the function of this gene during orofacial morphogenesis. However, while heterozygosity for this *Arhgap29* allele results in the absence of CL/P, it causes intra-oral adhesions, a well-accepted biomarker of cleft palate [41–43], providing evidence for a role, yet undefined, for ARHGAP29 in palatogenesis.

Prior whole mount and tissue *in situ* expression analyses show that the *Arhgap29* transcript is detected throughout embryonic craniofacial structures, including the palatal shelves [3]. Detailed immunofluorescence analysis revealed the presence of the ARHGAP29 protein in the mesenchyme, oral epithelium, and periderm of the palatal shelves, with particularly high expression in ectoderm-derived cells [3]. However, it is unclear which tissues require ARHGAP29 to promote proper palatogenesis and what the function of ARHGAP29 is during this process. This study is the first tissue-specific removal of ARHGAP29 used to investigate the mechanisms and requirement for ARHGAP29 during palatogenesis in the mouse. It reveals a novel role for ARHGAP29 in controlling the morphology, size and organization of epithelial cells covering the palatal shelves and preventing expression of contractility markers including phosphorylated myosin regulatory light chain (pMRLC) and alpha smoth muscle actin (α-SMA), in a spatial and temporal manner during palatogenesis.

## Results

### Generation and validation of *Arhgap29* conditional knockouts

ARHGAP29 is expressed throughout the oral cavity including the oral epithelium and periderm, both of which are ectoderm-derived. To evaluate the cell-specific requirements of *Arhgap29* during palatogenesis, we crossed an *Arhgap29* floxed allele [29] with either a *Tg(KRT14-cre)1Efu* (abbreviated *K14^Cre^)* [44] or an *Hhat>Tg(TFAP2A-cre)1Will* (abbreviated *Ect^Cre^)* [45, 46] driver to generate epithelial only, or ectoderm (epithelial and periderm) specific knockouts, respectively (Fig. 1A,B). The *Arhgap29* floxed allele contains LoxP sites flanking exons 4 and 5. These two exons are excised upon Cre recombination resulting in the production of a truncated nonfunctional ARHGAP29 protein that is targeted for degradation [39]. Additionally, we utilized a YFP reporter allele that is activated upon any Cre recombination (Fig. 1A,B). Immunofluorescence analysis of wild-type (WT) E14.5 embryos show strong expression of ARHGAP29 in ectoderm-derived tissues (oral epithelium and periderm) and concomitant absence of YFP in these same tissues (Fig. 1C-1C’’ and 1D-1D’’). ARHGAP29 is specifically reduced in only the basal epithelial cells of Arhgap29-K14^Cre+^ embryos, and not in the superficial periderm cells, while YFP is only detected in basal epithelial cells (Fig. 1E-1E’’ and 1F-1F’’). Conversely, we find a reduction of ARHGAP29 in all ectoderm-derived cells of Arhgap29-Ect^Cre+^ embryos, including in periderm cells (Fig. 1G-1G’’). Similarly, YFP is detected in both epithelial and periderm cells (Fig. 1H-1H’’). These data demonstrate the effective recombination of the *Arhgap29* floxed allele in the intended and distinct ectoderm-derived cell populations. Of note, all Arhgap29-Ect^Cre+^ mutants show some degree of YFP expression in the underlying mesenchyme (Fig. S1) suggesting some recombination of the *Arhgap29* floxed allele in mesenchymal cells either inherited from germline recombination or through ectopic Cre expression. Consequently, the Arhgap29-Ect^Cre^ should be considered a combined mesenchyme knockdown/ectoderm knockout rather than a strict ectoderm-specific conditional murine model, but will still be refered to as Arhgap29-Ect^Cre^ throughout our studies.

**Figure 1:**
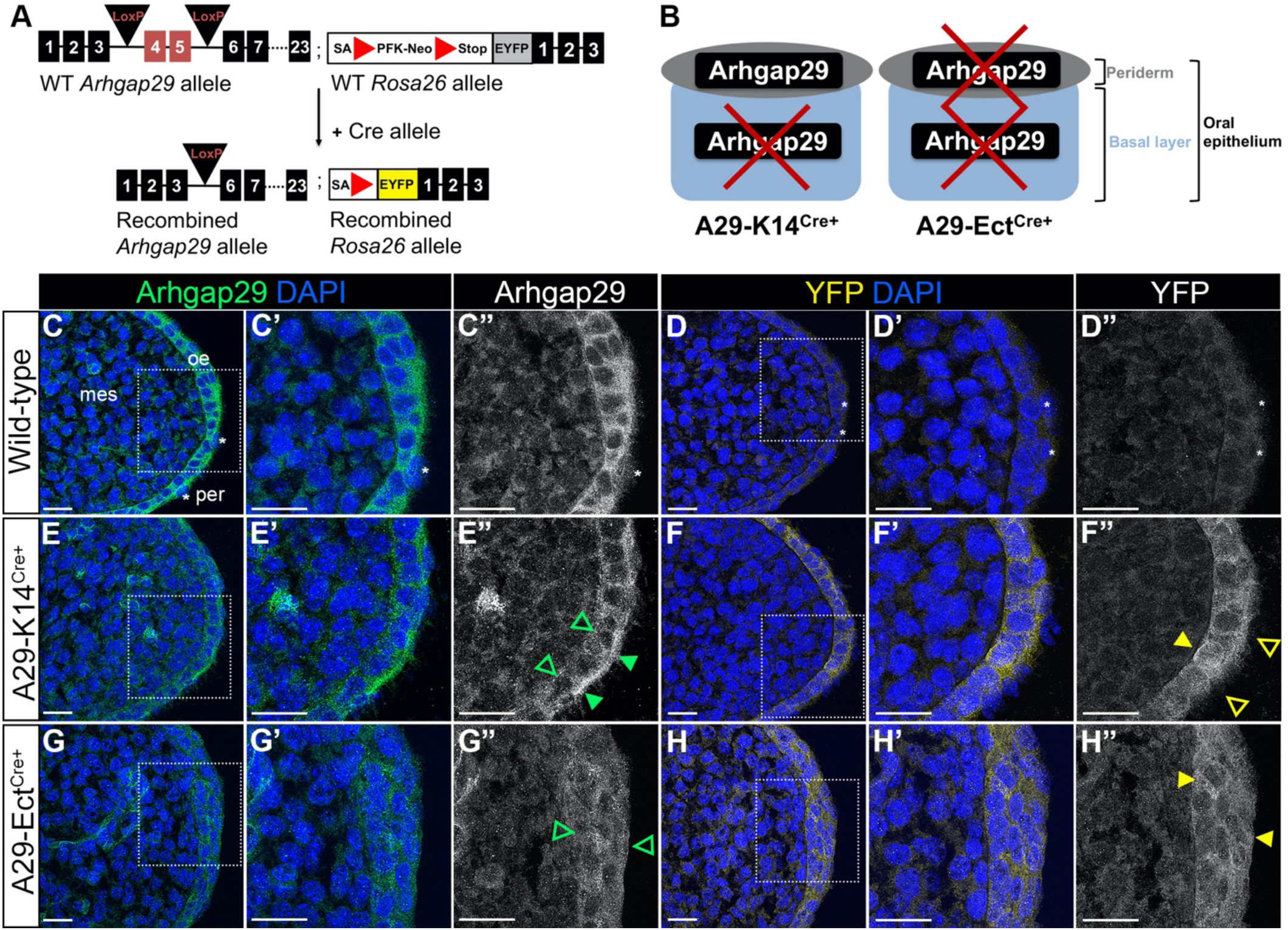
Validation of *Arhgap29* conditional knockouts. (A) Top, schematic of *Arhgap29* floxed allele and *Rosa26* allele harboring a YFP reporter. Bottom, upon Cre recombinase, the Lox P sites of the *Arhgap29* allele and the stop codons of the *Rosa26* allele are excised resulting in a truncated *Arhgap29* gene product and YFP expression, respectively. (B) Schematic of *Arhgap29* knockout in the Arhgap29-K14^Cre+^ (A29-K14^Cre+^; left) and Arhgap29-Ect^Cre+^ (A29-Ect^Cre+^; right) murine models. *Arhgap29* is removed only in basal epithelial cells in the A29-K14^Cre+^ and in the entire oral epithelium (basal and periderm cells) in the A29-Ect^Cre+^. (C-H”) Confocal slices of E14.5 elevated palatal shelf stained for ARHGAP29 (green in merged) and YFP (yellow in merged) of WT (C-D”), A29-K14^Cre+^(E-F”), and A29-Ect^Cre+^ (G-H”) embryos; boxed regions are shown at higher magnification on the right. Open or full arrow heads indicate cells lacking or expressing ARHGAP29 (green) or YFP (yellow), respectively. Asterisks denote periderm cells. mes = mesenchyme; oe = oral epithelium; per = periderm. Scale bars: 20 µm.

### Arhgap29-Ect^Cre+^ embryos display gross morphological defects

Full *Arhgap29* knockout animals are embryonic lethal, therefore we evaluated the genotypic frequencies of Arhgap29-K14^Cre+^ and Arhgap29-Ect^Cre+^ embryos at E14.5, E18.5, and post-weaning for viability. Arhgap29-K14^Cre+^ embryos exhibit the expected genotypic frequencies at each of these timepoints (Fig. 2A). Arhgap29-Ect^Cre+^ heterozygotes and homozygous embryos also display the expected genotypic frequencies at E14.5 and E18.5 (Fig. 2B). However, the weanling frequency of the homozygotes deviates from the expected mendelian ratio with a significant reduction in the number of mutants compared to littermate controls, suggesting a partial reduction in postnatal viability (Fig. 2B). Macroscopically, both WT and Arhgap29-K14^Cre+^embryos are morphologically indistinguishable at both E14.5 and E18.5 (Fig. 2C,D,G,H). However, Arhgap29-Ect^Cre+^ homozygous embryos at both E14.5 and E18.5 appear smaller compared to WT littermates and display a fully penetrant kinked tail (Fig. 2E,F,I,J). We identify occasional, yet significant Arhgap29-Ect^Cre+^ homozygotes with eye defects (Fig. S2A-E), syndactyly (data not shown), and general edema (data not shown). Adult Arhgap29-Ect^Cre+^ mice continue to exhibit a kinked tail of varying severity and a high frequency of “cataract-like” eye globe defects (Fig. S2D). At weaning, Arhgap29-Ect^Cre+^ homozygous pups exhibit a significant reduction in body weight, due to either weight loss or to an overall reduction in body size (Fig. S2F). Neither the Arhgap29-Ect^Cre+^ nor the Arhgap29-K14^Cre+^ animals displayed a cleft lip. Collectively, these results demonstrate that the combined loss of ARHGAP29 in the oral epithelium, periderm, and some mesenchymal cells, but not in the oral epithelium only, reduces embryonic viability and results in visible morphological defects, including a potential novel role for ARHGAP29 in eye development.

**Figure 2:**
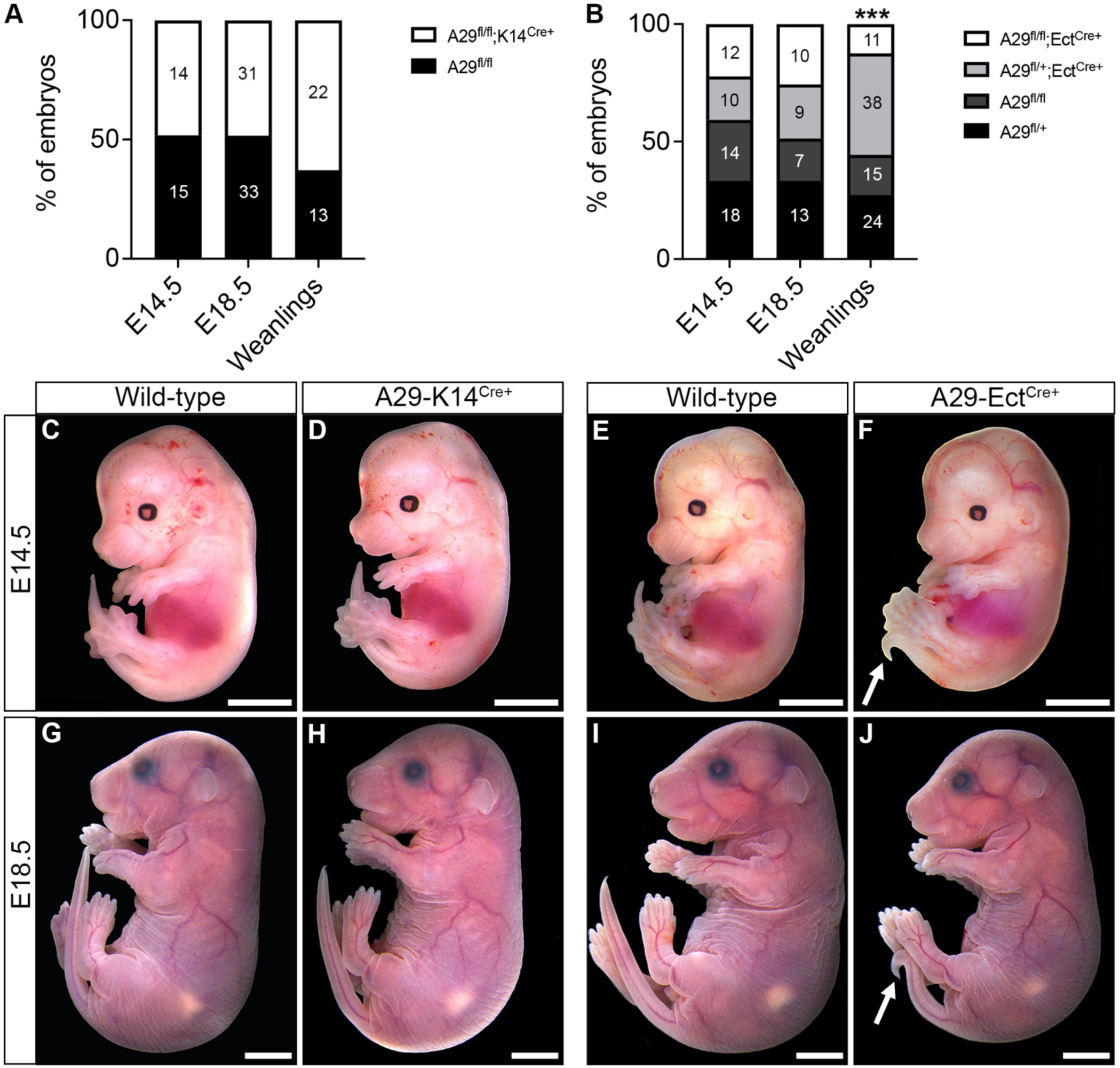
Loss of ARHGAP29 in ectoderm-derived, but not in basal epithelial cells only, leads to reduced survival and a kinked tail. (A-B) Genotypic frequencies expressed as percentage of embryos (raw numbers shown in each column) at E14.5, E18.5, and weanlings. \*\*\**P*<0.001, Pearson’s Chi-squared test. (A) A29^fl/fl^;K14^Cre+^ = A29-K14^Cre+^; A29^fl/fl^ = WT. (B) A29^fl/fl^;Ect^Cre+^ = A29-Ect^Cre+^; A29^fl/fl^ = WT. Note that A29^fl^;Ect^Cre+^ and A29^fl/+^ were part of the original cross but not further analyzed. (C-J) Macroscopic images of wild-type (C,G,E,I), A29-K14^Cre+^ (D,H), and A29-Ect^Cre+^ (F,J) embryos at E14.5 (C-F) and E18.5 (G-J).White arrows point to the kinked tail. Scale bars: 250 µm

### Arhgap29 is required in oral epithelial, periderm, and mesenchymal cells for proper palatogenesis

To determine if ARHGAP29 is required in ectoderm-derived cells, we performed histological analysis of coronal sections of Arhgap29-Ect^Cre+^ and control palates at E14.5 and E18.5. Palatal defects were scored in the anterior, middle, and posterior regions of the oral cavity, as either not elevated, elevated but not fused, or fused. At E14.5, Arhgap29-Ect^Cre+^ homozygous embryos display defects of the palate which includes palatal shelves that are not elevated, or elevated but not fused compared to WT (Fig. 3A-F). Interestingly, these defects are only significant in the anterior and posterior regions of the palate (Fig. 3G-I). To determine whether these defects are a simple delay in development or a permanent defect, we scored embryos for the presence of a cleft palate at E18.5. While all WT embryos display an intact palate at this timepoint, 23% of Arhgap29-Ect^Cre+^ homozygous embryos show the presence of a secondary cleft palate (Fig. 3J). Of the 5 embryos we observed with cleft palate, one displayed a partial cleft while four displayed a complete cleft. Given that Arhgap29-Ect^Cre+^ heterozygotes present with the expected genotypic frequencies without any gross morphological defects, delays in palatogenesis, or cleft palate (data not shown), we subsequently adapted our breeding scheme to only generate homozygous mutants.

**Figure 3:**
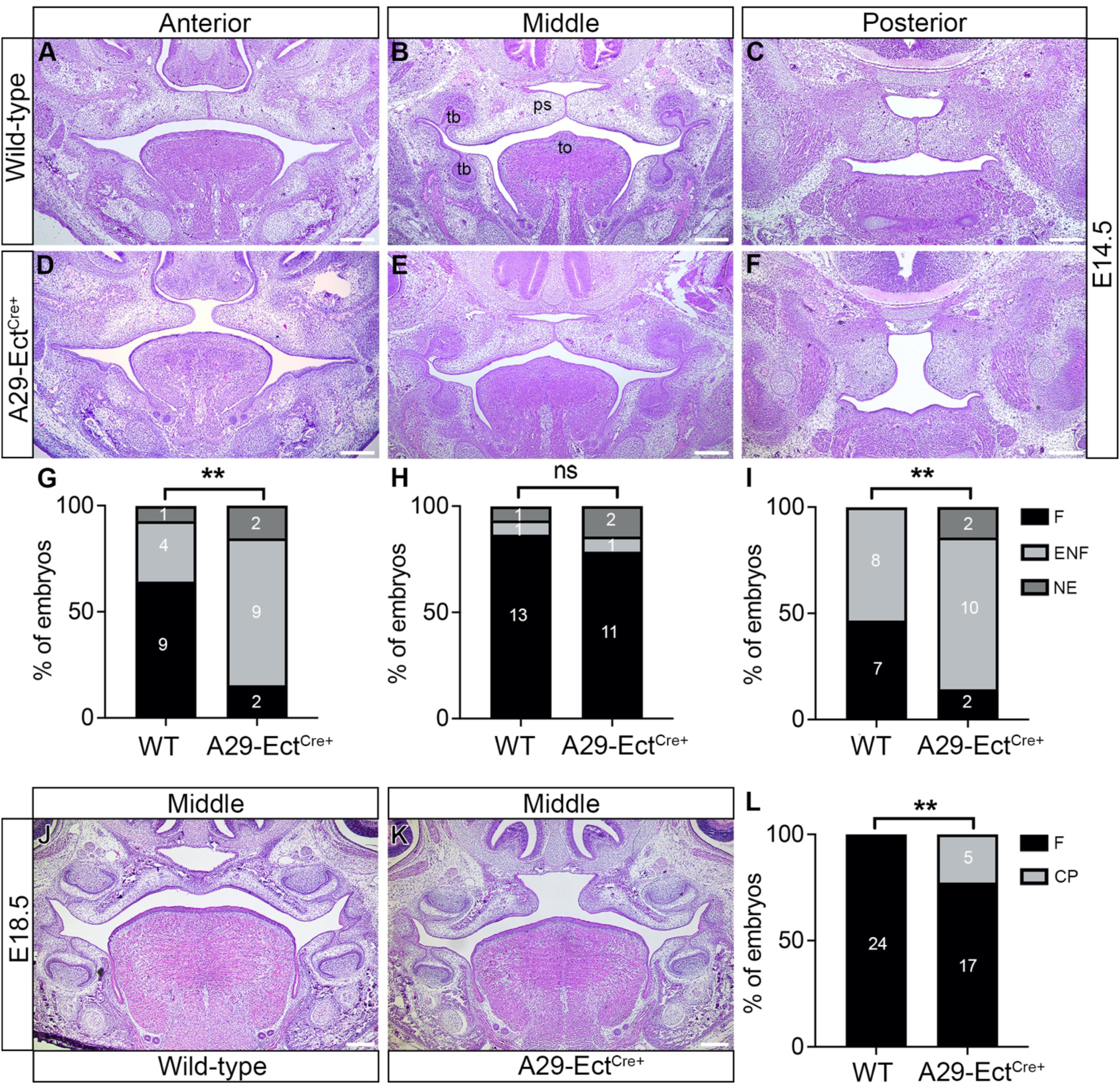
Arhgap29-Ect^Cre+^ embryos exhibit delayed palatogenesis and cleft palate. (A-F) Coronal sections of E14.5 wild-type (A-C) and A29-Ect^Cre+^ (D-F) embryos, stained with hematoxylin and eosin in the anterior (A,D), middle (B,E), and posterior (C,F) regions of the oral cavity. (G-I) Percentage of embryos (raw numbers shown in each column) with fused (F), elevated but not fused (ENF) or not elevated (NE) palatal shelves. (J-K) Coronal sections of E18.5 wild-type (J) and A29-Ect^Cre+^ (K) embryos, stained with hematoxylin and eosin in the middle region of the palate. (L) Percentage of embryos (raw numbers shown in each column) with fused (F) palatal shelves or cleft palate (CP). \*\**P* < 0.001, Fisher’s exact test; ns = not significant. ps = palatal shelf; to = tongue; tb = toothbud. Scale bars: 250 µm

Collectively, the analysis of Arhgap29-Ect^Cre+^ homozygotes demonstrates that ARHGAP29 is required in oral epithelial, periderm, and mesenchymal cells for proper palatogenesis.

### Specific loss of ARHGAP29 from oral epithelial cells does not disrupt palatogenesis

To determine if ARHGAP29 is required only in oral epithelial cells, we analysed the palatogenesis Arhgap29-K14^Cre+^, a model in which *Arhgap29* is deleted only in the oral epithelial cells. At E14.5, palatal shelves from both WT and Arhgap29-K14^Cre+^ embryos are adhered in the anterior, middle, and posterior palate (Fig. S3A-I). At E18.5, both WT and Arhgap29-K14^Cre+^ embryos display an intact palate that effectively separates the oral and nasal cavities (Fig. S3J,K). Together, because Arhgap29-Ect^Cre+^ show disrupted palatogenesis and Arhgap29-K14^Cre+^ do not, we conclude that ARHGAP29 is required in the ectoderm (oral epithelium and periderm) and the mesenchyme at E14.5 and E18.5 and that loss of ARHGAP29 from the basal oral epithelial cells alone is not sufficient to disrupt palatogenesis.

### ARHGAP29 is required for proper epithelial cell morphology of palatal shelves

While scoring Arhgap29-Ect^Cre+^ embryos for defects in palatogenesis, we observed a thickening of the epithelium covering the palatal shelves, specifically at their distal tip, the presumptive medial edge epithelium [47]. To quantify this observation, we measured the thickness of the epithelium along the shelves (when shelves down: lateral, tip, medial; when shelves up: lateral becomes oral, tip becomes MEE, medial becomes nasal) in the anterior, middle, and posterior regions of the palate in both embryos with palatal shelves not elevated and embryos with palatal shelves elevated but not fused. Fused shelves have varying degrees of medial edge epithelial cell-intercalation which precludes the measurement at this position. Arhgap29-Ect^Cre+^ embryos show a 1.5-2-fold increase in the thickness of the epithelium at the distal tip of the palatal shelves (Fig. 4A-L). This increase is restricted to the anterior region when the shelves are not elevated, and expanded to the entire antero-posterior regions when shelves are elevated (Fig. 4G-L). Additionally, after elevation, the oral and nasal epithelia of Arhgap29-Ect^Cre+^ embryos are significantly thicker in the anterior region of the oral cavity. Interestingly, as palatogenesis progresses and shelves elevate, the thickness of the epithelium increases all around the shelves in the anterior region, suggesting a worsening of the phenotype specifically anteriorly. Of note, the thickness of the epidermis around the head is unchanged between experimental groups (Fig. 4A-L) and all WT embryos display an elevated palate in the posterior region precluding the measurement of its epithelial thickness at that palatal position (Fig. 2I). Collectively, these results demonstrate a unique spatial and temporal requirement for ARHGAP29 in controlling epithelial morphology during palatogenesis.

**Figure 4:**
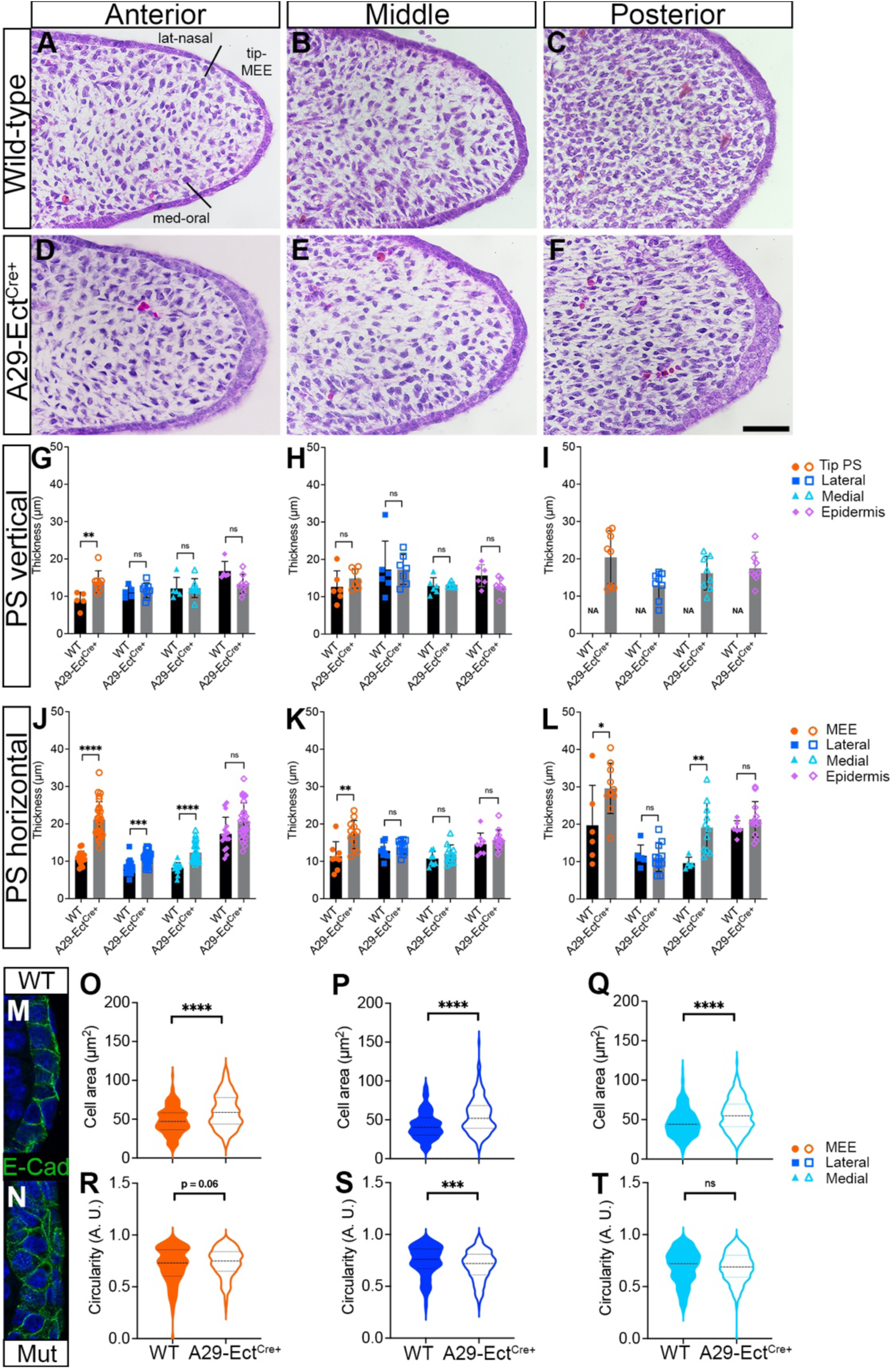
ARHGAP29 is required for oral epithelial cell morphology. (A-F) Elevated palatal shelves of E14.5 wild-type (A-C) and A29-Ect^Cre+^ (D-F) embryos, stained with hematoxylin and eosin in the anterior (A,D), middle (B,E), and posterior (C,F) regions of the oral cavity. Vertical black lines in (A) separate subregions of the palatal shelf epithelium used for measurements in panels G-T (lateral when vertical, nasal when horizontal; medial when vertical, oral when horizontal; tip of the palatal shelf when vertical, MEE when horizontal). Scale bars: 50 µm. (G-L) Quantification of epithelial thickness from WT and A29-Ect^Cre+^ E14.5 palatal shelves in a vertical (G-I) and horizontal (J-L) positions as well as thickness of the epidermis below the eye. (M-T) Using E-cadherin staining of WT and A29-Ect^Cre+^ [80] epithelial cells (examples shown in M and N), cell area (O-Q and circularity (R-T) were quantified. ****P < 0.0001, ***P<0.001, **P<0.01, ns = not significant, unpaired Student t-test. Error bars represent standard deviation.

### ARHGAP29 is not required for proliferation or cell death within the palatal shelves, but for epithelial cell morphology

Given the hypertrophic epithelium we observed in Arhgap29-Ect^Cre+^ mutants, we next investigated whether the thickening of the epithelium at the distal tip of the shelf is due to an increase in epithelial proliferation or changes in cell death. We focus on the anterior region of the oral cavity as this is where we observe the most significant change in epithelial thickness. The percentage of Ki-67-positive cells, the percentage of bromodeoxyuridine (BrdU)-positive cells 48 hours after its injection, and the percentage of TUNEL-positive cells are unchanged between WT and Arhgap29-Ect^Cre+^ mutants (Fig. S5). These data suggest that ARHGAP29 is not required for proliferation or cell death within the palatal shelves at E14.5. We then measured the size of the epithelial cells in this region and found that their area is larger in Arhgap29-Ect^Cre+^ compared to WT (Fig. 4O-Q), along with a reduction in their circularity only on the oral side of the shelves (Fig. 4R-T). Together, these data suggest that ARHGAP29 is not required for proliferation or cell death but regulates cell size and shape within the palatal shelves at E14.5.

### Arhgap29-Ect^Cre+^ embryos display defects in oral and lingual epithelial cell morphology and organization

To further explore causes of the hypertrophic palatal epithelium observed in Arhgap29-Ect^Cre+^, we then asked the question whether proteins required for epithelial development are properly expressed. Stratifin and keratin (K) 4, markers of early epithelial differentiation, are detected in the MEE [47] and oral epithelium adjacent to the tongue, respectively (Fig. 5A-C). No differences are observed between WT and Arhgap29-Ect^Cre+^ embryos (Fig. 5D-F), suggesting that premature differentiation is not contributing to the thickening of the palatal epithelium in these embryos. Next, we investigated the expression of E-cadherin, a cell-cell adhesion protein detected at the plasma membrane of epithelial cells, and K6, a marker of the periderm. In WT palatal shelves, the epithelial and periderm cells are each nicely organized into a single layer, with E-cadherin detected in all epithelial cells and K6 expression restricted to the periderm (Fig. 5G-H’). In Arhgap29-Ect^Cre+^, however, the epithelial and periderm cells are disorganized and are expanded into multiple layers (Fig. 5J), with K6 present in both basal and periderm cells (Fig. 5K,K’). Lastly, we evaluated the expression of annexin A1, a protein that facilitates the immune response with high expression in epithelial cells [48, 49]. In WT palatal shelves, annexin A1 is specific to and enriched in the periderm cells (Fig. 5I). Although the Arhgap29-Ect^Cre+^ palatal shelves also show the presence of annexin A1 in the periderm cells, the domain of expression is expanded to multiple superficial layers (Fig. 5L). Collectively, these data demonstrate a loss of specificity in spatial distribution of K6 and annexin A1, potentially revealing a lack of lineage specification. Changes in periderm function have been previously associated with changes in E-cadherin localization [50] and although our data show a similar pattern of expression in both WT and Arhgap29-Ect^Cre+^ palatal epithelium (Fig. 5G,J), E-cadherin staining reveals changes in epithelial shapes in A29-Ect^Cre+^ contributing to the disorganized epithelium. Interestingly, this disorganization is also observed in the lingual epithelium of Arhgap29-Ect^Cre+^ with a loss of classic rosette structures on the dorsum of the tongue, an increase in epithelial cell area and loss of circularity (Fig. S4).

**Figure 5:**
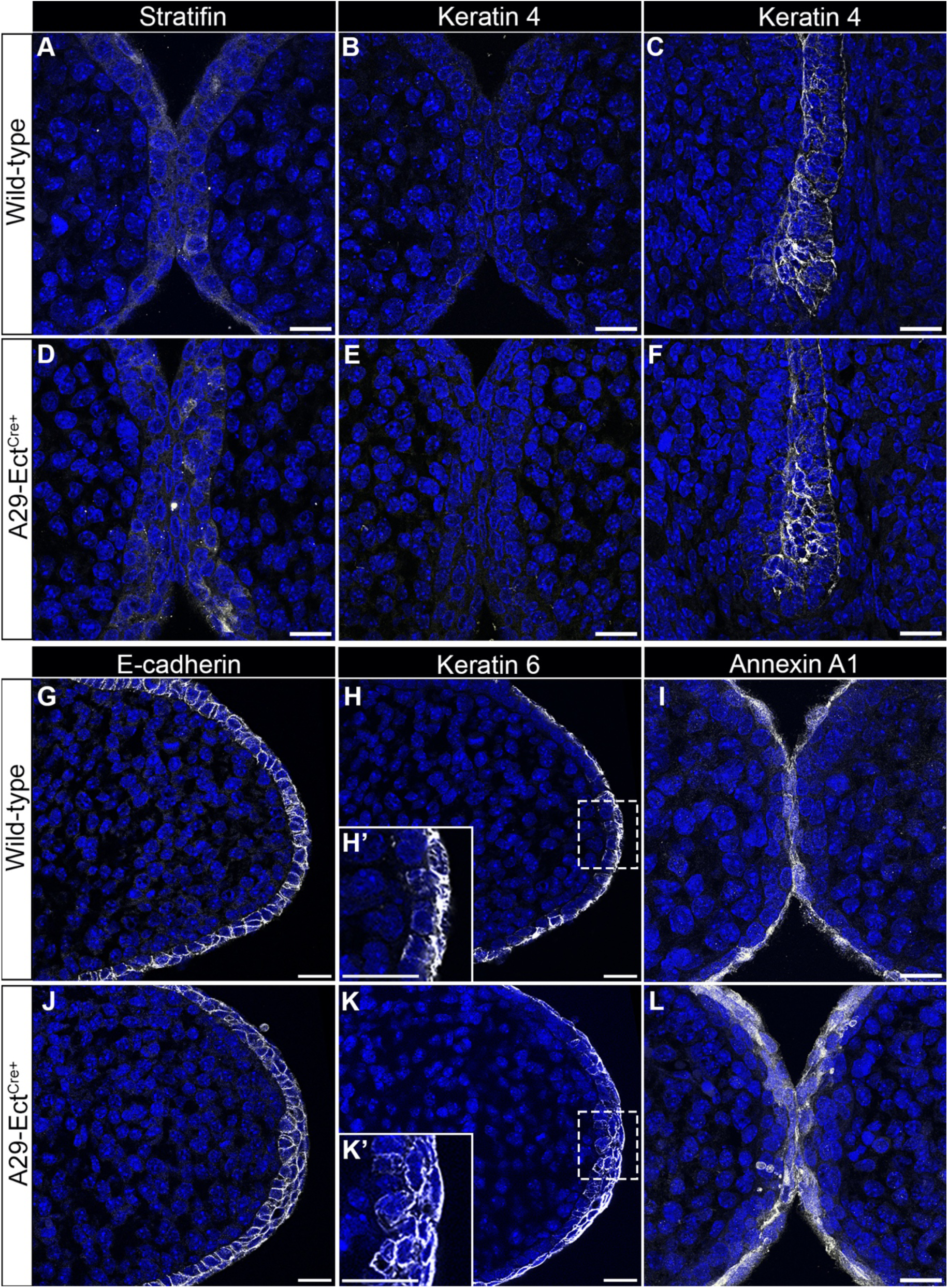
Periderm and epithelial organization, but not epithelial differentiation, are affected by ARHGAP29 loss. Confocal images of elevated palatal shelves and side of the tongue from E14.5 wild-type (A-C, G-I) and A29-Ect^Cre+^ (D-F, J-L) embryos stained with stratifin (A,B), Keratin 4 (B,C,E,F), E-cadherin (G,J), Keratin 6 (H,H’,K,K’) and Annexin A1 (I,L). (H’,K’) are higher magnification of the boxed regions of the same panel demonstrating changes in epithelial organization in A29-Ect^Cre+^. Scale bars: 20 µm

### Loss of ARHGAP29 promotes the expression of contractility markers in the MEE

Lastly, because ARHGAP29 is a regulator of the RhoA pathway, we asked whether the expression of RhoA downstream effectors is affected in Arhgap29-Ect^Cre+^ palatal epithelium. Myosin regulatory light chain (MRLC) is phosphorylated downstream of active RhoA and promotes contractility of the actin cytoskeleton [51]. In WT palate epithelial cells, p-MRLC is present at low levels in basal cells and enriched at cell-cell junctions of superficial cell layers, sites of presumptive mechanical tension (Fig. 6A). In Arhgap29-Ect^Cre+^, however, p-MRLC is upregulated throughout the epithelium, including strong signal in the superficial layers (Fig. 6B). To further demonstrate a change in contractility, we immunostained palatal shelves with alpha-smooth muscle actin (α−SMA), another downstream target of RhoA typically detected in activated myofibroblasts and only present in specialized epithelial cells [52–54]. Our data confirm its absence in WT palatal epithelium, but show a strong upregulation in Arhgap29-Ect^Cre+^ palatal epithelium during fusion (Fig. 6A-B). These results demonstrate that in the absence of ARHGAP29, contractility markers are upregulated, which may contribute to the morphological changes in the oral and lingual epithelium and could perturb cytoskeletal remodeling required for on-time palatogenesis.

**Figure 6:**
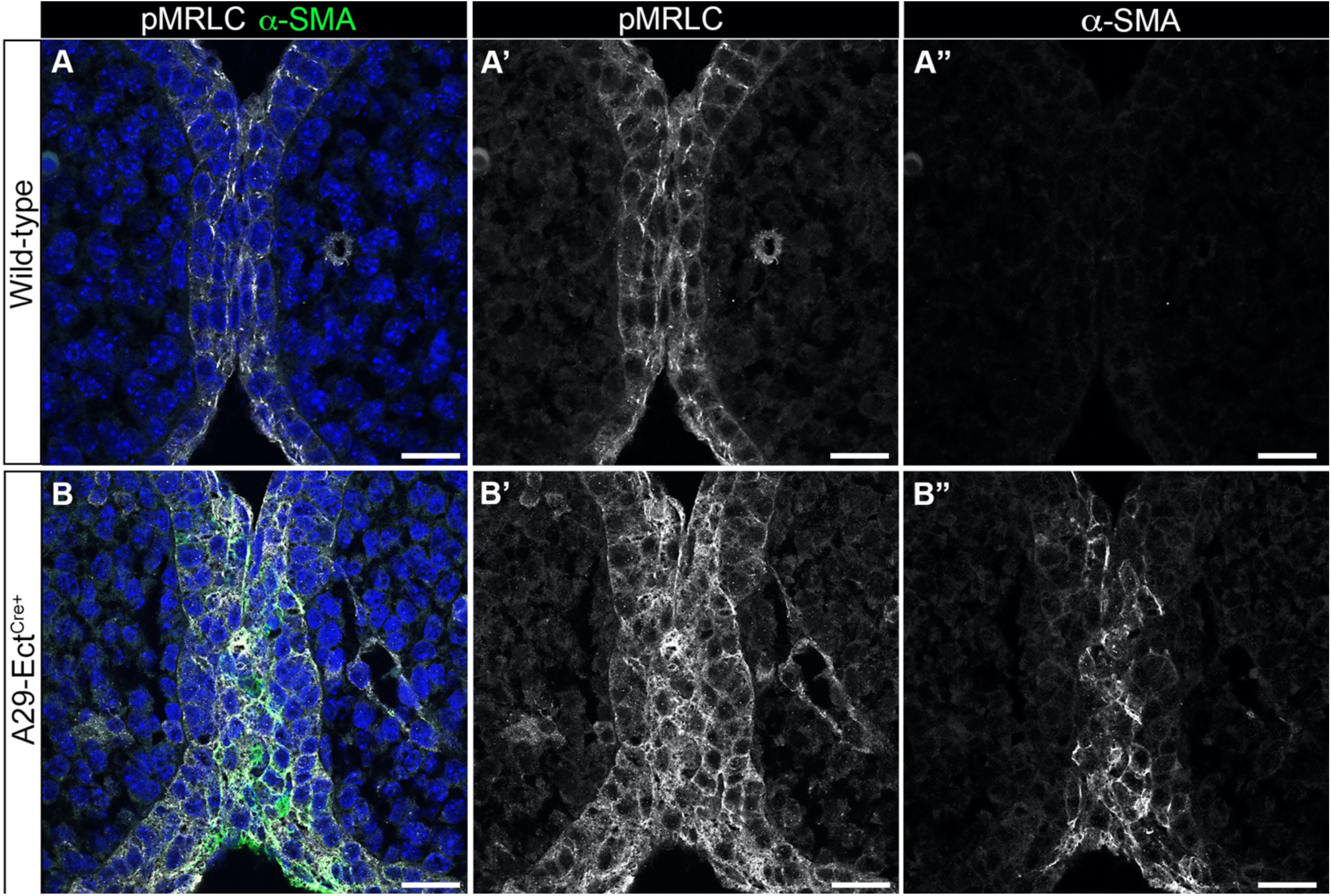
Loss of ARHGAP29 in oral epithelial and periderm cells increases contractility protein expression. Confocal images of elevated palatal shelves from E14.5 wild-type (A-A”) and A29-Ect^Cre+^ (B-B”) embryos dual stained with phosphor-myosin regulatory light chain (pMRLC, white in merged) and alpha smooth muscle actin (α-SMA, green in merged). Scale bars: 20 µm

## Discussion

In this study, we used tissue-specific deletion of *Arhgap29* in mice to provide the first direct evidence that ARHGAP29 contributes to proper palatal elevation and fusion. Our results show that loss of ARHGAP29 in ectoderm- and mesenchyme-derived cells of the developing face results in a significant delay in secondary palatal fusion at E14.5, and a significant yet not fully penetrant cleft palate at E18.5. Because our studies preclude us to follow palatogenesis of individual embryos as they develop, we cannot ascertain that an embryo with palatal shelves down at E14.5 would have a cleft palate at E18.5. However, it is tempting to speculate that the 21% of Arhgap29-Ect^Cre+^ embryos with palatal shelves not elevated or fused at E14.5 would be the ones with the most delayed palatogenesis ultimately leading to the 23% of E18.5 embryos with a cleft palate. And despite only about 20% of embryos exhibiting a cleft palate, this incomplete penetrance is frequently observed in murine models in which genes causing human clefting are deleted, such as *Afadin*, *Specc1L* or *Grhl3* [20, 23, 55, 56].

ARHGAP29 belongs to the RhoA pathway, which has been previously implicated in OFC [3, 57], yet how it may contribute to palatogenesis has been unclear because of a lack of useful tools, including our previous *Arhgap29* loss-of-function allele due to early lethality [38]. Immediately downstream of ARHGAP29 and RhoA is the ROCK/myosin signaling pathway. Activation of this pathway results in increased filamentous actin and phosphorylation of myosin regulatory light chain (MRLC). Dynamic changes in the actomyosin cytoskeleton are known to be required for proper palatogenesis [58–64]. Additionally, murine mutants for *β-catenin*, *Specc1l,* and *Nectins (Nectin-1* and *Nectin-4),* modulators of the actomyosin cytoskeleton, display cleft palate [20, 24, 25, 65]. Consistent with these studies, we found that loss of *Arhgap29* early during facial development leads to increased expression of phosphorylated MRLC in oral epithelial cells, suggesting that altered actomyosin dynamics contributes to the mechanism by which ARHGAP29 promotes palatogenesis (Fig. 7). These observations are further supported by our previous *in vitro* studies showing that cutaneous keratinocytes knockeddown for ARHGAP29 display increased filamentous actin and pMRLC compared to control [34]. Along those lines, we also detected the presence of alpha-smooth muscle actin (αSMA) in epithelial cells at the tip of the palatal shelves in Arhgap29-Ect^Cre+^ embryos. αSMA is expressed mostly in fibroblasts when they become contractile but is also detected in some epithelial cell populations with contractile properties like in submucosal glands of the airway [66]. Outside of one report detailing the presence αSMA in the oral epithelium of fusing palatal shelves in rat [67], there is currently no data to suggest a role for this protein in normal palatogenesis. We believe that the presence of αSMA in conjunction with an increase in pMLRC could increase the overall contractility of epithelial cells of the palatal shelves, rendering them stiffer and less amenable to remodeling during palatogenesis (Fig. 7). Direct measurements of tissue stiffness or actomyosin dynamics *in vivo* would need to be performed to formally test such a mechanism.

**Figure 7:**
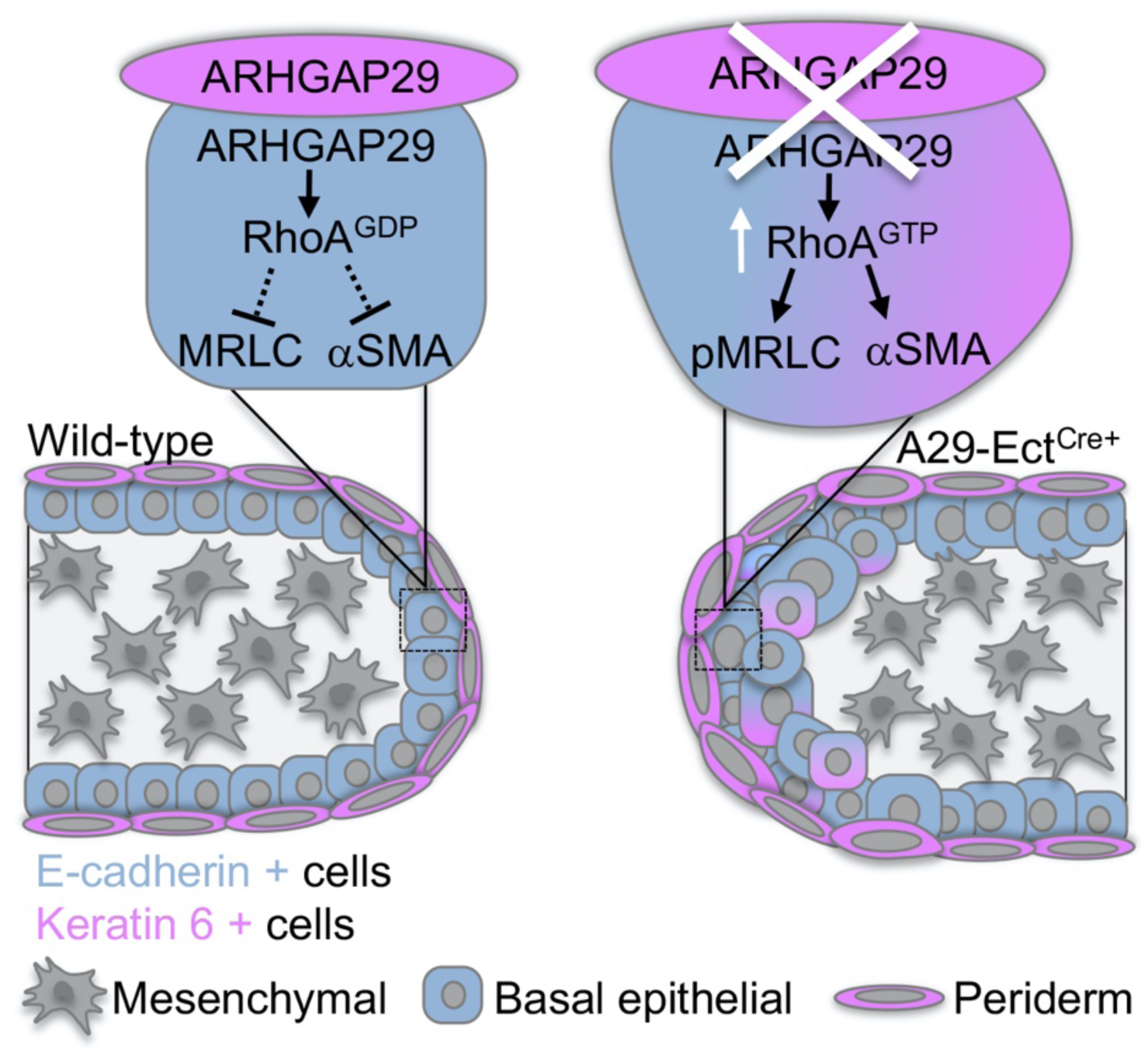
Proposed model for the role of ARHGAP29 during palatogenesis. Left, ARHGAP29 within the basal epithelium, and periderm of wild-type PS, functions to inactivate RhoA which limits the expression of pMRLC and αSMA. Right, loss of ARHGAP29 in the basal epithelial and periderm cells of A29-Ect^Cre+^ embryos results in increased epithelial thickness, increased epithelial cell size, and defects in epithelial organization. Loss of ARHGAP29 results in the increase in active RhoA, which promotes phosphorylation of myosin regulatory light chain (pMRLC) and alpha smooth muscle actin (α-SMA). Collectively, these defects in epithelial organization, morphology, and contractility may perturb the dynamic remodeling of the palatal shelves required for their elevation and fusion.

Our detailed observations reveal epithelial hypertrophy in Arhgap29-Ect^Cre+^ mutants, particularly at the tip of the palatal shelves. This phenotype is reminiscent of the oral epithelial thickening observed in the *Tbx1* knockout mouse [68]. However, contrary to what we observe in our mutants, the thickening in the *Tbx1* knockout embryos results from a decrease in proliferation and abnormal epithelial differentiation. Epithelial hypertrophy has also been reported in palatal organ cultures treated with nocodazole, a drug that disrupt microtubules [69]. Interestingly, this defect does not implicate cell proliferation, but involves altered cytoplasmic localization of GEF-H1, a microtubule-associated Rho GEF that couples microtubule dynamics to cell contractility. Interplay between the RhoA-ROCK-actomyosin pathway and microtubules is well established [70] and microtubule dynamics has been shown previously to be involved in palatogenesis [23, 71], although there is currently no evidence of a role for ARHGAP29 in regulating microtubule dynamics. Epithelial thickening without changes in proliferation is also evident in the epidermis of mice conditionally lacking laminin 332 in keratinocytes [72]. In these mice, αSMA is strongly upregulated in the superficial epidermal layers, and the shape of epidermal cells is altered. This is consistent with our findings, where loss of ARHGAP29 leads to larger epithelial cells around the palatal shelves with reduced oral epithelial circularity. This increase in size likely accounts for the hypertrophic epithelium observed in these mutants, and is consistent with significant increase in cell size and alterted morphology in ARHGAP29-knockdown keratinocytes cultured *in vitro*. [34]. Cell-cell and cell-matrix adhesions often influence cell shape and size and ARHGAP29 has been previously shown to be involved in endothelial cell adhesion dynamics during tubulogenesis [39]. Whether ARHGAP29 contributes to similar adhesion remodeling during palatal shelf elevation is currently unknown.

Although palatal shelves appear as homogenous outgrowths from the maxillary processes in the oral cavity, molecular heterogeneity along the anterior-posterior axis is well established [11]. Our results extend this heterogeneity to cellular morphology as the oral epithelial hypertrophy in Arhgap29-Ect^Cre+^ mutant is initially only detected at the tip of the palatal shelves when they are vertical, and then expanded to the medial and lateral epithelia covering the shelves following their elevation. Even more noticeable is that the defect is restricted to the anterior region of the oral cavity when the shelves are vertical and then extends to the posterior region after their elevation. How palatal shelves change from a vertical to a horizontal position is still unclear, yet Coleman [73] and Walker [74] previously proposed a heterogeneous mechanism along the anterior-posterior axis, whereby the anterior part of the palatal shelves elevate by a “flip-up” process, while the posterior and middle parts of the palatal shelves elevate by a remodeling mechanism [73–75]. It is also clear that physical forces are involved [76], yet how they contribute to the changes in palatal shelf positions are not well understood. Because ARHGAP29 regulates the actomyosin cytoskeleton, it is tempting to speculate that it may contribute to force generation required for changing palatal shelf position, and may do so heterogeneously along the anterior-posterior axis, further supporting different mechanisms for palatal shelf elevation.

Given our previous studies on ARHGAP29 and single-cell RNA sequencing data sets of embryonic facial structures, we knew ARHGAP29 is predominantly detected in ectoderm-derived cells [3, 34, 38, 77]. This ectoderm enrichment is why we used Cre-drivers that would target epithelial cells. Our results using the K14-Cre and the Ect-Cre drivers demonstrate that ARHGAP29 in basal oral epithelial cells alone is not required for palatogenesis. Its removal from all ectoderm-derived cells (oral basal and periderm), however, impairs palatogenesis and disrupts other developmental processes (i.e., kinked tail, eye defect, lower body weight). Given that we observe partial recombination outside of ectoderm-derived cells using this driver, we cannot conclude whether this defect is due solely to the loss of ARHGAP29 in ectoderm-derived cells or to its potential reduction in other tissues. The use of mesenchymal specific or neural-crest specific drivers followed by a comparison with our current models would help tease apart the real contribution of ARHGAP29 to these different embryonic phenotypes.

We previously reported a loss-of-function *Arghap29* allele and showed the presence of oral tissue adhesions in heterozygous embryos [38]. We originally postulated that this phenotype was due to the reduction of ARHGAP29 in the periderm, and that despite periderm cells being present in these embryos, somehow the reduction of ARHGAP29 was impairing periderm function. To our surprise, we did not observe oral adhesions in either the Arhgap29-K14^Cre+^ or Arhgap29-Ect^Cre+^ heterozygotes or homozygotes (data not shown). The lack of oral adhesions in the Arhgap29-K14^Cre+^ can likely be explained by the fact that ARHGAP29 is still present in the periderm, however, their absence in the Arhgap29-Ect^Cre+^ are more puzzling. Because the loss-of-function heterozygous model led to a 50% reduction of ARHGAP29 from time of conception, and the changes in ARHGAP29 levels in Arhgap29-Ect^Cre+^ mutants occurred mid-gestation, one may speculate that there may be a non-cell autonomous temporal requirement for ARHGAP29 function in the periderm. Additionally, our previous loss-of-function heterozygous model was in a full C57BL6/J background, while the Arhgap2929-Ect^Cre+^ has a mixed genetic background, suggesting that the more complex genetic background of the Ect-Cre driver may also contribute to this divergent phenotype.

In conclusion, by properly organizing epithelial cells around the palatal shelves and modulating the expression of contractility proteins, ARHGAP29 may participate in mechanotransduction that play a critical role in time-sensitive embryonic tissue dynamics. Learning how ARHGAP29 integrates with other signaling pathways will further our understanding of palatal shelves movement during normal development and pathogenic orofacial clefts.

## Methods

### Generation and processing of experimental mouse lines

All animal procedures were approved by the Animal Care and Use Committee of the University of Iowa. All mice utilized in this study, except Ect^Cre^, were of C57BL/6J background. Female mice homozygous for the FLAG tagged floxed *Arhgap29* allele (Arhgap29^fl/fl^; generated at the University of Iowa Core Facility) and for the YFP Rosa-26 allele (ROSA^yfp/yfp^; Jax strain #006148) were bred with A29^fl/fl^; ROSA^yfp/yfp^;Cre-positive males, harboring either Keratin-14 Cre (Tg(KRT14-cre)1Efu) or *Ect^Cre^* (Tg(Tcfap2a-cre)1Wil) alleles. *K14^Cre^*is driven by the Keratin-14 promoter and fully activated by E11.75 in basal epithelial cells [44] while Ect^Cre^ is driven by an TFAP2a upstream enhancer element and is active at E8.5 in all ectoderm-derived cells [45]. The presence of a copulatory plug is designated as embryonic (E) 0.5. Females were injected with bromodeoxyuridine (BrdU, 5 mg/g body weight) intraperitoneally two hours prior to euthanasia then euthanized with CO2. Litters were harvested at E14.5 and E18.5 as previously described [38]. Heads were dissected and fixed in 4% paraformaldehyde and tail snips or yolk sac used for genotyping. Heads were further processed and embedded in paraffin.

### Histological analysis and phenotype scoring

Serial coronal paraffin sections (7µm thick) were obtained and stained with hematoxylin and eosin for histologic analysis. Images were acquired using a Nikon Eclipse E-600 (Nikon, Melville NY) and measurement of palatal epithelium thickness was performed using Nikon NIS-Element software. Although embryos were harversted 14 days after male and female were put together in a cage, serial sectioning revealed phenotypic variations amongst litters as judged by standardized criteria of E14.5 craniofacial development (palatal shelves elevated and opposite epithelia in contact). To provide consistency in our analyses, we applied the same filtering criteria we previously developed [38]. Briefly, we only considered the following litters for our analysis: (1) they contained at least one WT to be used as internal reference and (2) at least one of these WT demonstrated elevated and fused palatal shelves. Using these criteria, we analyzed 16 litters of Arhgap29-K14^Cre+^ and 9 litters of Arhgap29-Ect^Cre+^ at E14.5. This filtering strategy was not necessary at later timepoints. Palatal shelf positions were scored as “not elevated” (vertical on the side of the tongue), “elevated but not fused” (horizontal above the tongue with a gap in the midline between them), and “fused” (shelves in contact with medial edge epithelium or medial edge seam still present). Oral cavity/palatal regions were defined as anterior (immediately anterior to toothbuds), middle (eye and toothbuds in the field), and posterior (presence of medial pterygoid processes).

### Fluorescent immunostaining and microscopy

Immunostaining of paraffin sections was performed as previously described [78]. Four percent PFA fixed, paraffin embedded slides were de-paraffinized, permeabilized with 0.5% TritonX-PBS and then treated with 0.05% Tweeen-20 citric acid antigen retrieval. Samples were blocked with 2% normal goat serum (Vector Laboratories, Burlingame, CA) in 1% PBS-Bovine serum albumin overnight at 4°C, stained in primary antibody at 37°C for 2 hours and stained with secondary antibody for 1 hour at 37°C. Nuclear DNA was labeled with Hoechst 33342 (1/10,000). Samples were mounted using ProLong^TM^ Diamond Antifade Mountant. Images were acquired with a Zeiss 980 confocal microscope (Zeiss, Oberkochen, Germany) and processed using Fiji software [79].

### Proliferation analysis via BrdU and Ki67 detection

BrdU and Ki67 were detected following immunofluorescent staining as previous described [38]. Tiled confocal images of the oral cavity were acquired and processed using Fiji software. Percentage of proliferating cells was calculated by dividing the number of BrdU+ or Ki67+ cells by the total number of cells as identified by positive nuclear Hoechst staining. Palatal shelves were defined by drawing a verticle line from the palatal hinge region on the nasal side to the oral side of the shelf. Cells were counted in the palatal shelf epithelium and mesenchyme.

### Epithelial cell morphology analysis

Immunofluorescent staining was performed on E14.5 coronal sections as described above. Embryos with elevated but not fused palatal shelves were stained for E-cadherin and Hoechst 33342 to label the cell membrane and nuclear DNA, respectively. Confocal images of the oral cavity were acquired and processed using Fiji software. Individual cells were traced using the E-cadherin staining, and cell area and circularity calculated using the measure command in the Fiji software. Palatal shelves were defined as described in proliferation analysis section.

### Antibodies

List of all antibodies used in the study is provided in supplemental Table 1.

## Acknowledgements

The authors are grateful to all the MarTina lab members for support, and to Dr. Eric Van Otterloo for the use of his microscope and TUNEL reagents. The authors acknowledge funding support from grants R01AR067739 (to MD), F31DE033222 (to EA) and T32GM145441 (to EA) from the National Institute of Health; from the Graduate Program Student Government Research Grant (to EA); from the Iowa Center for Research by Undergrauates (CM); and from the Department of Anatomy and Cell Biology. The authors also gratefully acknowledge the support of the Central Microscopy Research Facility and the Transgenic Core Facility of the University of Iowa. The authors declare no competing financial interests.

## Author contributions

Emily Adelizzi: Conceptualization, Data curation, Formal analysis, Writing the first draft and review-editing.

Lindsey Rhea: Conceptualization, Formal analysis, Writing – review and editing

Campbell Mitvalsky: Data curation Samuel Pek: Data curation Bethany Doolittle: Data curation.

Martine Dunnwald: Conceptualization, Data curation, Formal analysis, Funding requisition, Supervision, Writing – review and editing.

