## Supplemental figures for "The ectodermal loss of ARHGAP29 alters epithelial morphology and organization and disrupts murine palatal development"

**Table S1. List of antibodies used in the study.**

| Antibody Name | Host Species | Company Name | Company Location | Catalog # | Dilution |
|---|---|---|---|---|---|
| **Primary Antibodies** | | | | | |
| GFP | Rabbit Polyclonal | Invitrogen | Waltham, MA | A6455 | 1/400 |
| Keratin 4 | Rabbit Polyclonal | Invitrogen | Waltham, MA | MA537810 | 1/3200 |
| Annexin A1 | Rabbit Polyclonal | Invitrogen | Waltham, MA | 713400 | 1/5000 |
| Ki67 | Rabbit Monoclonal | Invitrogen | Waltham, MA | RM-9106-S | 1/200 |
| Stratifin | Goat Polyclonal | Santa Cruz | Dallas, TX | SC-7683 | 1/100 |
| E-Cadherin | Mouse Monoclonal | BD Biosciences | San Jose, CA | 610181 | 1/100 |
| Arhgap29 | Rabbit Polyclonal | Proteintech | Rosemont, IL | 12583-1-AP | 1/100 |
| Keratin 6 | Rabbit Polyclonal | Covance | Princeton, NJ | PRB-169P | 1/500 |
| Bromodeoxyuridine | Rat Monoclonal | Abcam | Cambridge, UK | ab6326 | 1/100 |
| Alpha-Smooth Muscle Actin | Mouse Monoclonal | Sigma-Aldrich | St. Louis. MO | A2547 | 1/8000 |
| Phosphorylated-Myosin Regulatory Light Chain | Rabbit Polyclonal | Cell Signaling | Danvers, MA | 3671 | 1/100 |
| **Secondary Antibodies** | | | | | |
| Alexa Fluor 488 goat anti-mouse | Mouse | Invitrogen | Waltham, MA | A-11001 | 1/400 |
| Alexa Fluor 488 goat anti-rabbit | Rabbit | | | A-11008 | 1/400 |
| Alexa Fluor 568 goat anti-mouse | Mouse | | | A-11004 | 1/400 |
| Alexa Fluor 568 goat anti-rabbit | Rabbit | | | A-11011 | 1/400 |
| Anti-Rat IgG-FITC | Rat | Chemicon | Rolling Meadows, IL | AP136F | 1/200 |

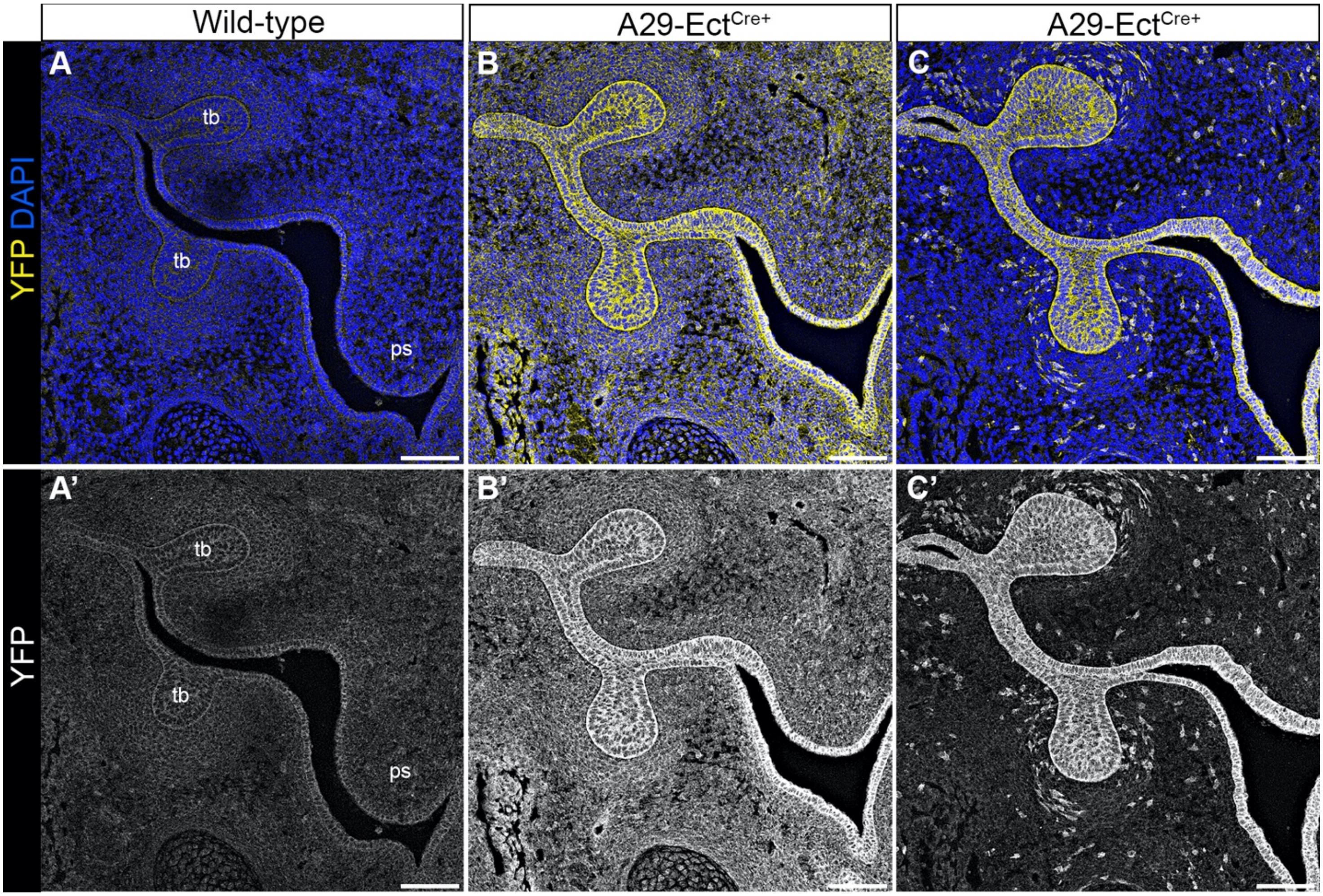


**<u>Supplemental Figure 1:</u> Characterization of Ect$^{Cre}$ expression pattern.** (A-C') Confocal images of the oral cavity (coronal sections) of E14.5 wild-type (A,A') and A29-Ect$^{Cre+}$ (B-C') embryos, stained with YFP (yellow in merged). ps = palatal shelf; tb = toothbud. Scale bars: 25 μm

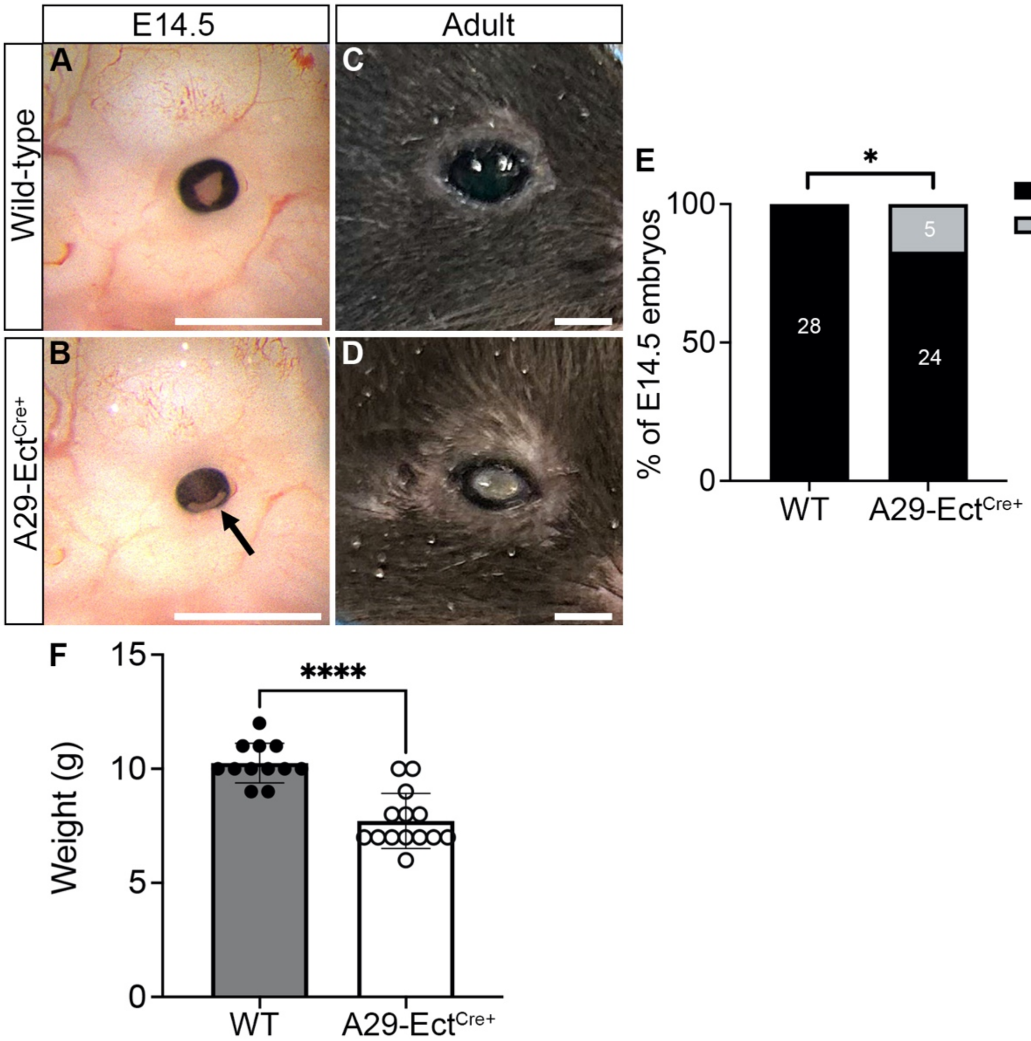


**<u>Supplemental Figure 2:</u> Loss of ARHGAP29 leads to ocular defects and weight loss.** (A-D) Macroscopic images of E14.5 (A,B) and adult (C,D) WT (A,C) and A29-Ect$^{Cre+}$ (B,D) eye. Black arrow indicates ocular defect. Scale bars: 200 μm. (E) Quantification of eye defects in E14.5 embryos. (F) Quantification of weanling body weight. ****$P$ < 0.0001, unpaired Student t-test; *$P$ < 0.05, Chi-square test.

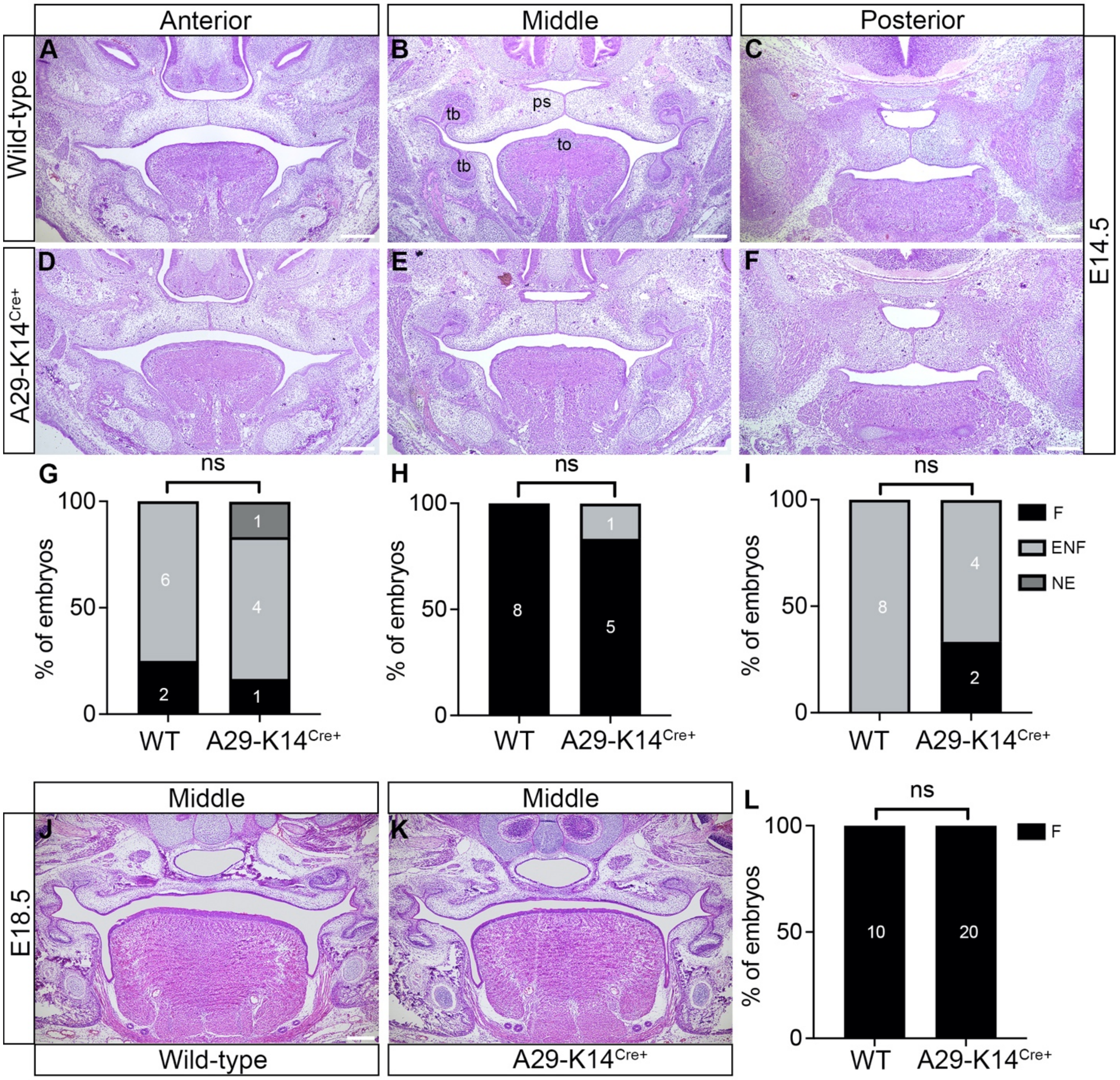


**Supplemental Figure 3: ARHGAP29 in oral epithelial cells only is not required for proper palatogenesis.**

(A-F) Coronal sections of E14.5 wild-type (A-C) and A29-K14$^{Cre+}$ (D-F) embryos, stained with hematoxylin and eosin in the anterior (A), middle (B), and posterior (C) regions of the oral cavity. (G-I) Percentage of embryos (raw numbers shown in each column) with fused (F), elevated but not fused (ENF) or not elevated (NE) palatal shelves. (J-K) Coronal sections of E18.5 wild-type (J) and A29-K14$^{Cre+}$ (K) embryos, stained with hematoxylin and eosin in the middle region of the palate. (L) Percentage of embryos (raw numbers shown in each column) with fused (F) palatal shelves or cleft palate (CP). ns = not significant, Fisher's exact test. ps = palatal shelf; to = tongue; tb = toothbud. Scale bars: 250 µm

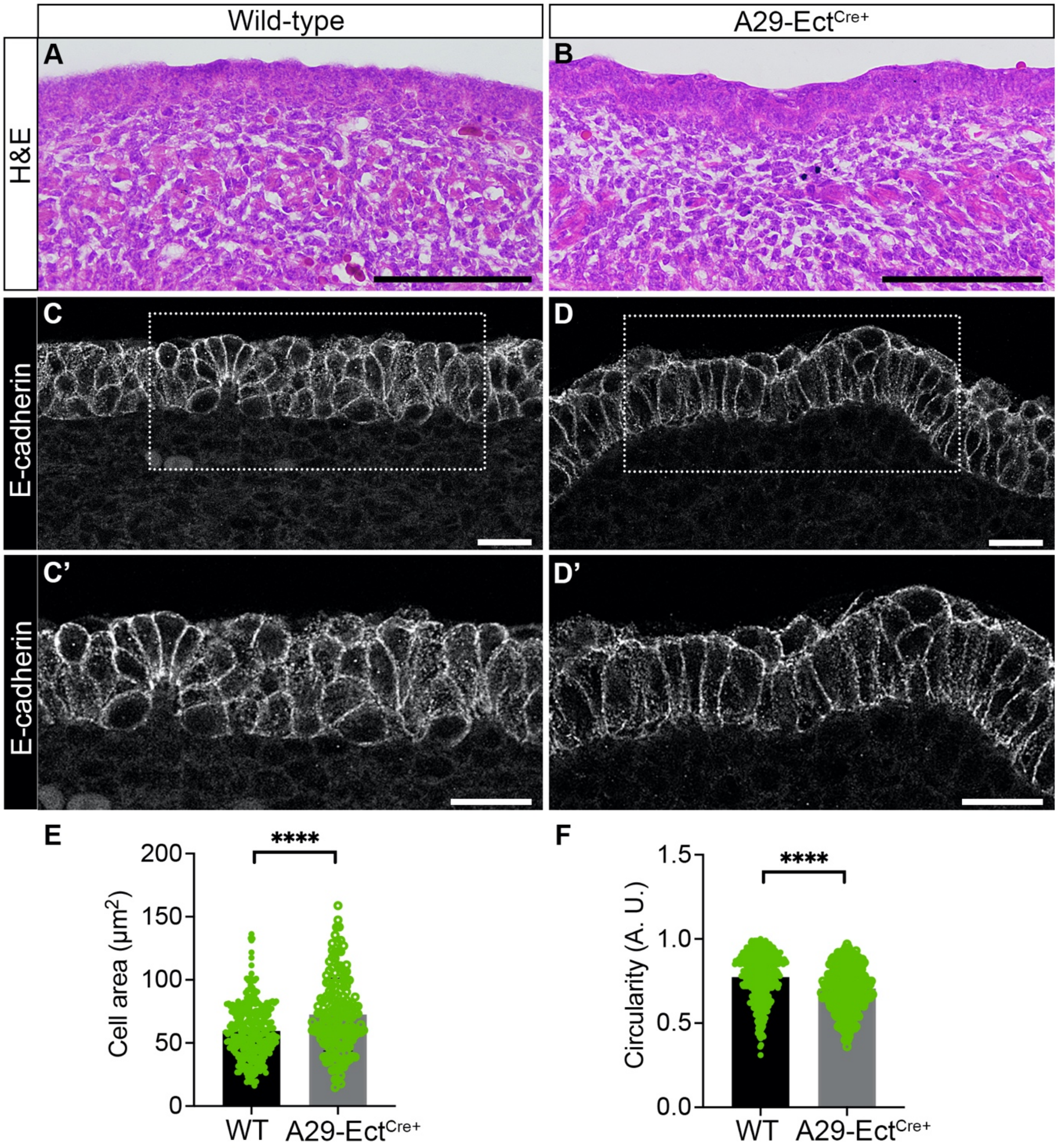


**Supplemental Figure 4: A29-Ect$^{Cre+}$ embryos exhibit a disorganized lingual epithelium.** (A,B) Coronal section of E14.5 WT (A) and A29-Ect$^{Cre+}$ (B) tongue stained with Hematoxylin and Eosin. (C-D') Confocal images (single slice) of lingual coronal sections of E14.5 WT (C,C') and A29-Ect$^{Cre+}$ (D,D') stained with E-cadherin. Magnified images of panels C and D are shown in panels C' and D', respectively. Scale bars: 20 µm. (E) Quantification of lingual epithelial cells area. (F) Quantification of lingual epithelial cell circularity. ****$P$ < 0.0001, unpaired Student t-test.

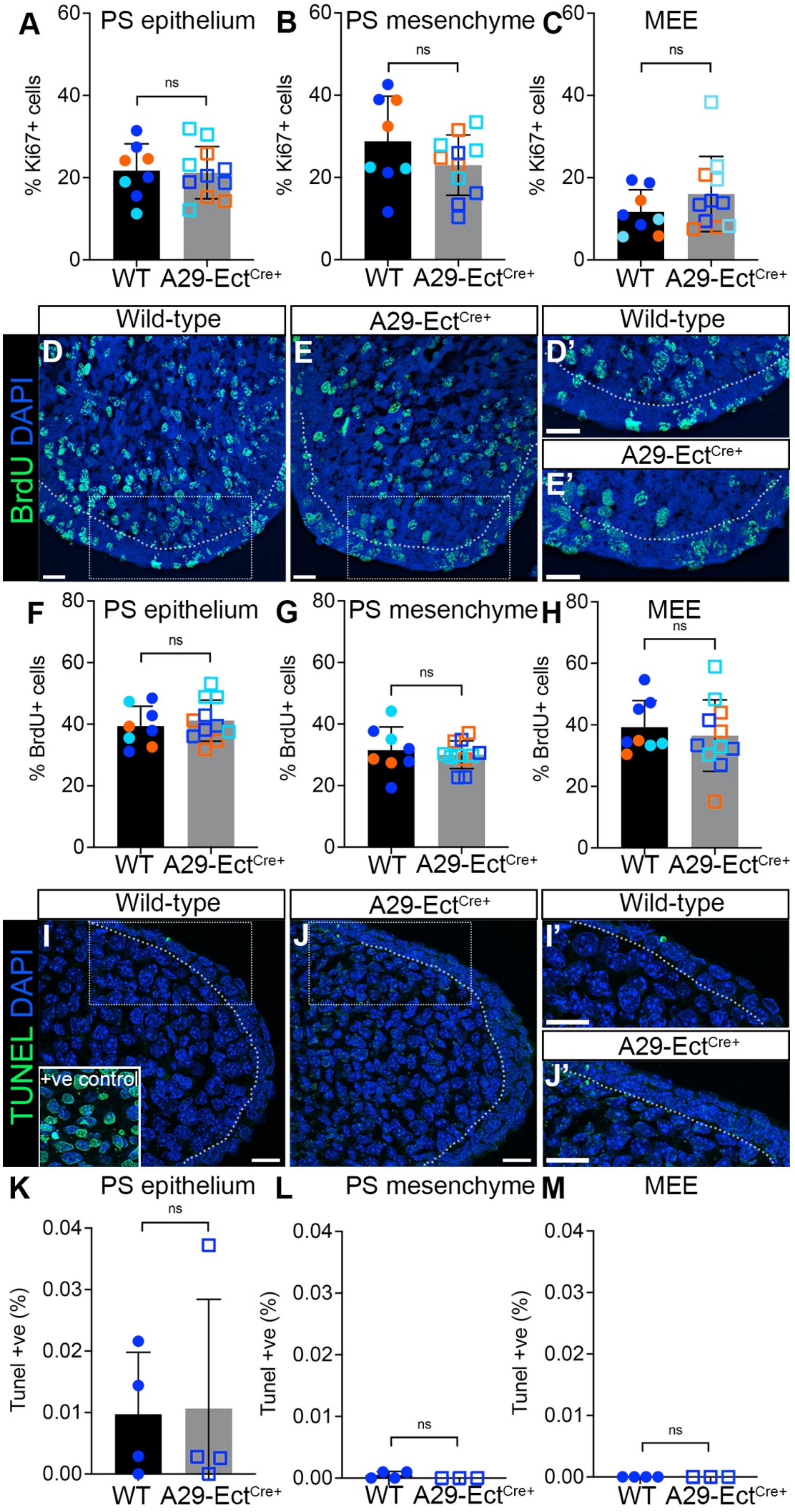

A
PS epithelium
% Ki67+ cells
ns
B
PS mesenchyme
C
MEE
WT
A29-Ect$^{Cre+}$
D
E
D'
E'
Wild-type
A29-Ect$^{Cre+}$
BrdU DAPI
F
G
H
% BrdU+ cells
I
J
I'
J'
TUNEL DAPI
+ve control
K
L
M
Tunel +ve (%)

**<u>Supplemental Figure 5:</u> Arhgap29 is not required for prolifeation or apoptosis during palatogenesis.** (A-C) Quantification of the percentage of Ki67 positive cells in E14.5 palatal shelf (PS) epithelium (A), mesenchyme (B), and medial edge epithelium (MEE, C). (D-E') Coronal sections of E14.5 WT (D,D') and A29-Ect$^{Cre+}$ (E,E') palatal shelves stained with BrdU (green). Magnified images of panels D and E are shown in panels D' and E', respectively. (F-H) Quantification of the percentage of BrdU positive cells in E14.5 PS epithelium (F), mesenchyme (G), and MEE (H). (I-J') Coronal sections of E14.5 WT (I,I') and A29-Ect$^{Cre+}$ (J,J') palatal shelves stained with TUNEL (green). Magnified images of panels I and J are shown in panels I' and J', respectively. (K-L) Quantification of the percentage of TUNEL positive cells in E14.5 PS epithelium (K), mesenchyme (L), and MEE (M). Dotted white lines indicate the basement membrane. Symbols correspond to different palatal shelf positions (orange = fused; cyan = not elevated; blue = elevated but not fused). ns = not significant, unpaired Student t-test. Scale bars: 20 μm
